# GORASPs link Golgi cisternae laterally to stabilize the rims and prevent vesiculation

**DOI:** 10.1101/2020.04.28.066381

**Authors:** Rianne Grond, Tineke Veenendaal, Juan Duran, Ishier Raote, Sebastiaan Corstjens, Laura Delfgou, Benaissa El Haddouti, Vivek Malhotra, Catherine Rabouille

## Abstract

In vitro experiments have shown GRASP65 (GORASP1) and GRASP55 (GORASP2) proteins function in stacking Golgi cisternae. However, in vivo depletion of GORASPs in metazoans have given equivocal results. We have generated a mouse lacking both GORASPs and find that Golgi cisternae remained stacked. However, the stacks are disconnected laterally from each other and the cisternal cross-sectional diameters are significantly reduced compared to their normal counterparts. These data support earlier findings on the role of GORASPs in linking stacks and we suggest that unlinking of stacks affects dynamic control of COPI budding and vesicle fusion at the rims. The net result is that cisternal cores remain stacked, but cisternal diameter is reduced by rim consumption.

## Introduction

Most eukaryotic cells exhibit stacked Golgi cisternae. What holds the cisternae together and what is the function of cisternal stacking? Identification of stacking factors could help address these questions.

### Discovery of GRASP proteins in cisternal stacking

In mammalian cells, pericentriolar stacks of Golgi cisternae are separated from each other late in G2 and, by metaphase, these membranes are converted into tubules and small vesicles. Cisternal stacks are visible in telophase and post cytokinesis, when they return to the pericentriolar region of the newly formed cells (Lucocq & Warren, 1987; Pecot & Malhotra, 2006). Warren and colleagues reconstituted the fragmentation of isolated Golgi stacks using mitotic cytosol (Misteli & Warren, 1995) and the reverse process, assembly of Golgi stacks from mitotic Golgi fragments (Rabouille, Misteli, Watson, & Warren, 1995). Treatment of the disassembled Golgi membranes with an alkylating agent (N-ethyl maleimide or NEM) inhibited stack reassembly (Rabouille et al., 1995). This led to identification of a **G**olgi **R**e**a**ssembly **S**tacking **P**rotein 65, GRASP65(Barr, Puype, Vandekerckhove, & Warren, 1997) (now called GORASP1) that is N-terminally myristoylated. This modification, along with tight binding to the coiled-coil Golgi protein GM130(Nakamura, Lowe, Levine, Rabouille, & Warren, 1997) are required for attachment of GORASP1 to cis- and medial-Golgi cisternae (Bachert & Linstedt, 2010). Remarkably, adding antibodies targeted against GORASP1, or a recombinant GORASP1 to disassembled Golgi membranes, prevented assembly of cisternae into stacks (Barr et al., 1997; Wang, Seemann, Pypaert, Shorter, & Warren, 2003). GRASP55(GORASP2), a protein highly homologous to the N-terminal half of GORASP1, localizes to more distal Golgi cisternae and was also subsequently shown to be required for stacking in vitro (Shorter et al., 1999). Moreover, simultaneous depletion of both GORASPs by siRNA in HeLa cells, resulted in unstacked Golgi profiles, consistent with small single cisternal and Golgi fragments (Wang, Satoh, & Warren, 2005; Xiang & Wang, 2010). These data led to the conclusion that GORASPs are essential to stack Golgi cisternae (Rabouille & Linstedt, 2016; Vinke, Grieve, & Rabouille, 2011).

### Concerns about the role of GORASPs in cisternal stacking

Deletion of the single GORASP gene expressed in Pichia pastoris, (Levi, Bhattacharyya, Strack, Austin, & Glick, 2010) and Drosophila cells (Kondylis, Spoorendonk, & Rabouille, 2005) is largely inconsequential with respect to Golgi cisternal stacking. siRNA-based depletion of GORASP1 in HeLa cells did not affect the overall Golgi stacking, but the numbers of cisternae/stack were reduced. Interestingly, these cells enter mitosis, but do not proceed beyond mitosis because of defects in spindle dynamics (Sutterlin, Polishchuk, Pecot, & Malhotra, 2005). In a transgenic murine system, deletion of GORASP1 did not affect animal viability, or the stacking of cisternae, but the stacks were laterally separated from each other (Veenendaal et al., 2014). This phenocopies earlier reports on the functional significance of GORASP1 and 2 in connecting cisternae laterally without a requirement in the cisternal stacking (Feinstein & Linstedt, 2008; Jarvela & Linstedt, 2014; Puthenveedu, Bachert, Puri, Lanni, & Linstedt, 2006). However, these studies addressed the effect of depleting either only one of the two GORASP orthologs. In the mouse knockout of GORASP1 that failed to detect an obvious effect on Golgi stacking (Veenendaal et al., 2014), the levels of GORASP2 were unaffected, and given that GORASP2 is the more abundant of the two GORASPs, the possibility of a functional redundancy could not be excluded. To address redundancy, Wang and colleagues used CRISPR–Cas9 to completely deplete GORASPs in HeLa and HEK293 cells, but these alterations led to a significant reduction (up to 75-80% of the control levels) of GM130 and Golgin 45 proteins (Bekier et al., 2017). Fluorescence microscopy-based analysis with antibodies to GM130 and the late Golgi specific, transmembrane protein TGN46, revealed that these membrane components were dispersed throughout the cytoplasm instead of localized at the Golgi as in control cells. Not clear, therefore, was whether this was due to loss of GORASPs, loss of 75-80% of GM130, Golgin 45, or perhaps loss of another cellular component that might be depleted by these procedures (see Fig 4.,(Bekier et al., 2017)). But even more concerning are the electron micrographs shown in that paper, which show the presence of either disorganized, stacked Golgi membranes, or single isolated cisternae that could well be ER (Figure 5D). In sum, there is a general agreement that loss of a single GORASP protein (by siRNA, CRISPR or gene knockout) in mammalian cells or its orthologous gene in flies or *Pichia pastoris* has no effect on cisternal stacking. The results from the double depletion of GORASPs in HeLa and HEK293 cells are far less clear.

### A potential resolution to the functional significance of GORASPs

To address the apparent discrepancy regarding the role of GORASPs in Golgi cisternae stacking, we generated a mouse in which both GORASPs are knocked out. The general physiology of the mouse is under investigation, but development is delayed compared to the control and mice are therefore smaller and lighter in weight. Also, analyses of the small intestine during early post-natal development as well as intestinal organoids reveal that GORASPs are not required to stack Golgi cisternae. The Golgi stacks are separated from each other as reported previously (Feinstein & Linstedt, 2008; Veenendaal et al., 2014), but more striking is the finding that the cisternal diameter is significantly reduced in diameter compared to control Golgi cisternae.

We discuss our findings to help resolve debates on the role of GORASPs in stacking of Golgi cisternae.

## Results and Discussion

### Quantitation of GORASP levels in the mouse DKO

Standard procedures to obtain a double GORASP1 and GORASP2 knockout failed because of embryonic lethality. We therefore employed an alternative, albeit routine, procedure to obtain a mouse lacking GORASP1 and GORASP2.

A GORASP1 mouse knockout (Veenendaal et al., 2014) was crossed with a conditional knock-out (cFlox) mice, expressing endogenous GORASP2, with exon 5 flanked by loxPsites (tm1 c allele). Crossed with Rosa Cre ERT2, tamoxifen induces Cre recombinase, under control of Zp3 promotor, eliminating GORASP2 exon 5 expression resulting in a mouse lacking both functional GORASP1 and GORASP2. Following injection with Tamoxifen (TAM), the intestine was isolated for further analysis. In parallel, from uninjected mice, we generated small intestine budding organoids that were then incubated with 4-hydroxy TAM (4 OT) to deplete GORASP2(see Methods section). These tissues were first analyzed for the levels of GORASP1 and GORASP2. As expected, these tissues were completely depleted of GORASP1. Our procedure for depletion of GORASP2 resulted in excision of exon 5 from the genomic locus (Figure 1C, D), whereas exons 1-4 and 6-10 are still present. This yields an exon 5-lacking mRNA (truncated by 131 bp) that is expressed at 20-30% of the levels seen in the control *(Supp fig. S1A-C).* We checked whether this mRNA produces a protein product. The small intestine of TAM and oil-injected mice were dissected at Postnatal (P) day 10, P21 and P42 and processed for immunofluorescence (IF) using a GORASP2 antibody raised against its C-terminus (see Methods section). We failed to detect GORASP2 protein in any of the TAM injected mice (**Figure 2A for P21, *Supp fig. S2 for P10***), while it was abundant in the control animals. We also processed the treated organoids and found a complete loss of full-length GORASP2 protein expression as shown by immunofluorescence (Figure 2B) and by Western blot (Figure 2C). The data suggest that deletion of exon 5 either destabilizes GORASP2 mRNA, or results in a base shift, yielding a protein with an aberrant C-terminal sequence. The level of this truncated mRNA is reduced to 20-30% when compared with non-treated controls (*Supp fig. S1C*), thus it appears to be largely degraded. Furthermore, if it were to harbor a base shift, it would be translated into a protein harboring PDZ1, half of PDZ2 and a very short C-terminus due to the presence of stopcodons in exon 6. In either case, we would not detect a full length protein in these animals.

**Figure 1:**
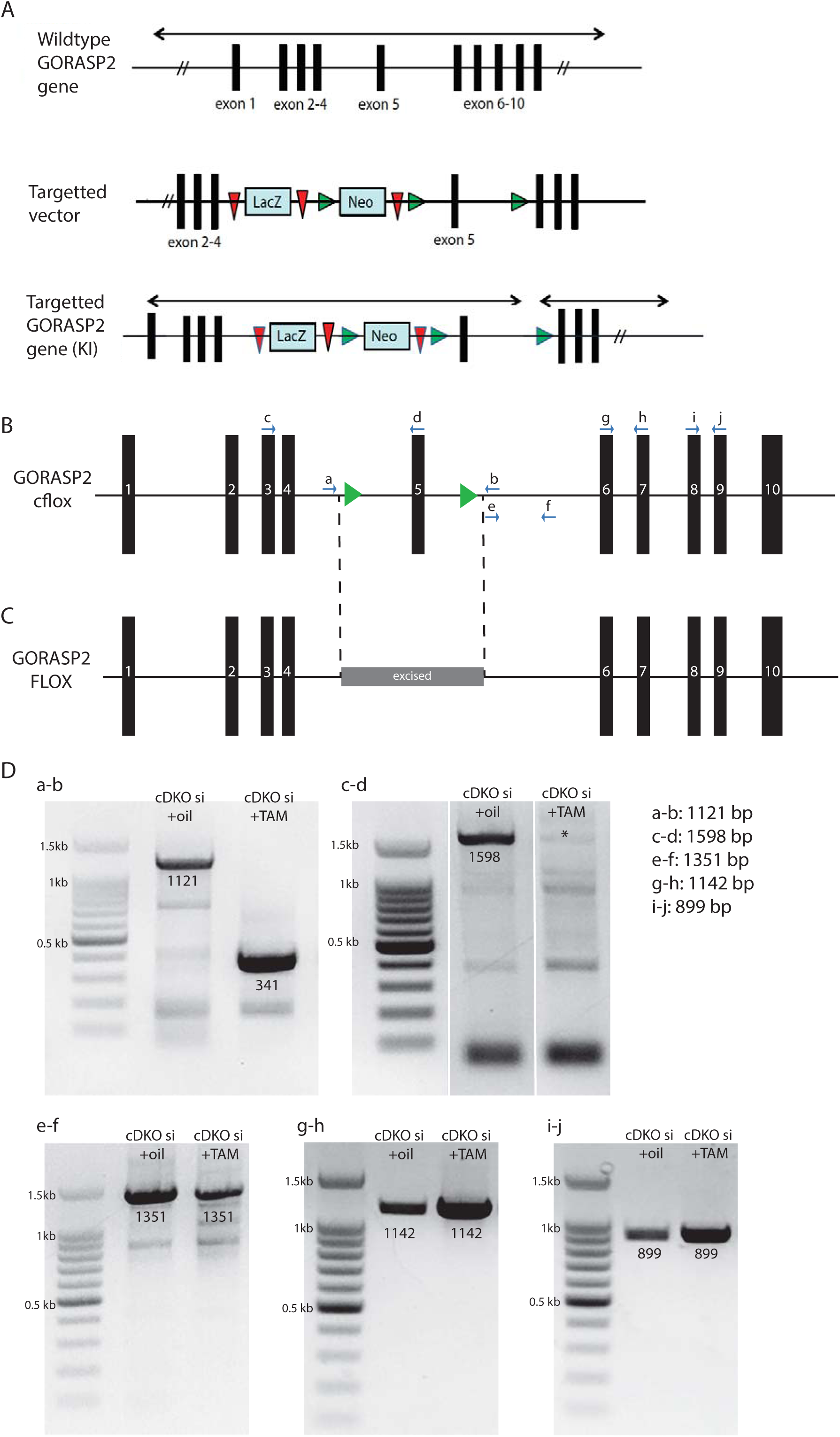
GORASP KI, cFlox and FLOX allele. **A:** Schematic of the GORASP2 KI allele with the LacZ/Neo cassette between exons 4 and 5. **B:** Schematic of the GORASP2 cFlox allele upon crossing GORASPKI with Flp-recombinase expressing mice. The exon 5 is now flanked by two LOXP sites. **C:** Schematic of the GORASP2 FLOX allele upon TAM injection (after crossing with a Rosa Cre ERT2 mouse). The exon 5 is excised. **D:** A series of PCR using different combination of primers showing that Exon5 is deleted (a-b and c-d), that a part of intron 5 is still present (e-f), that exons 6 and 7(g-h) and exons 8-9(i-j) are still present.

**Figure 2:**
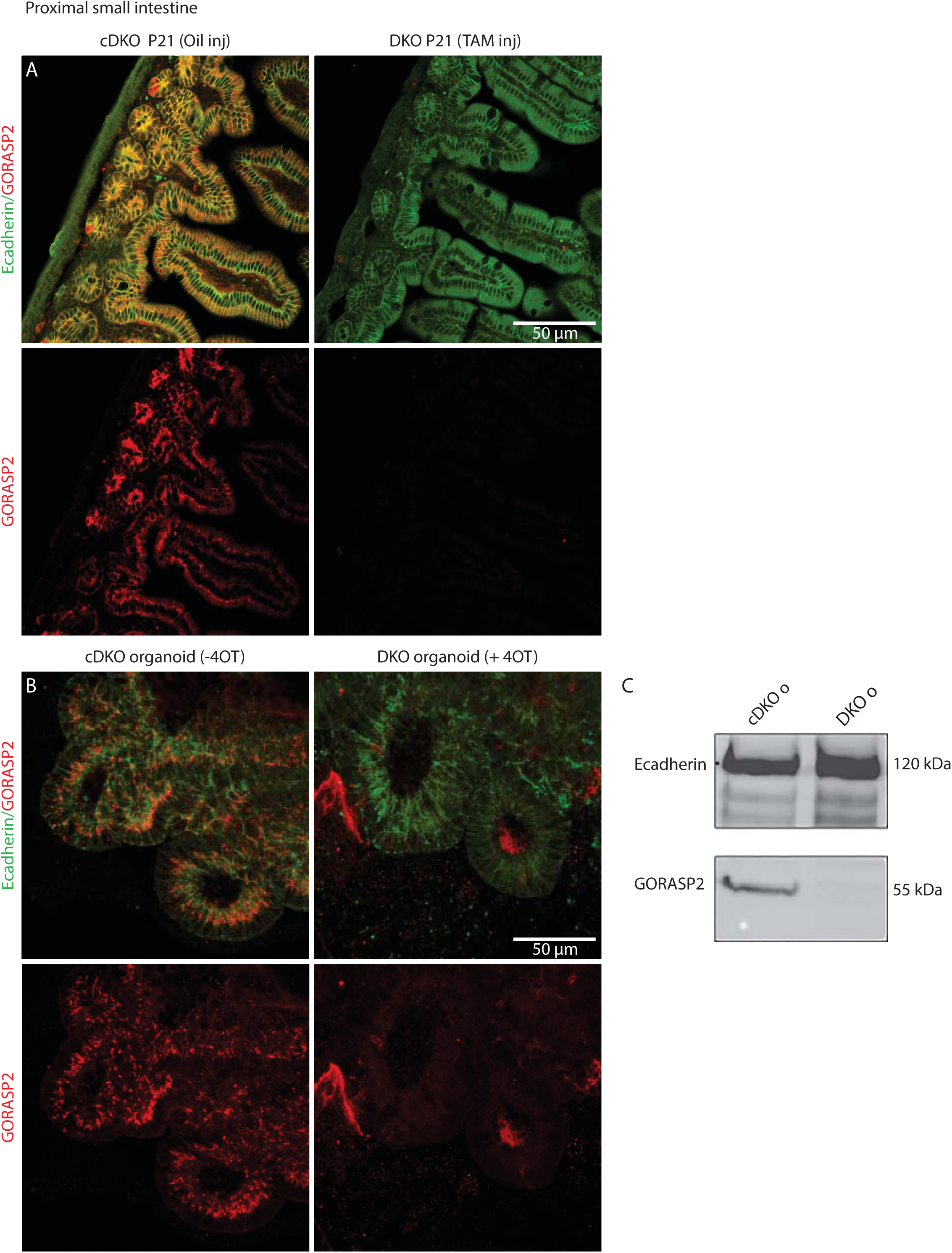
GORASP2 protein is no longer expressed in mouse small intestine and organoids upon TAM/4 OT exposure. **A:** Visualization of GORASP2 (red) and E-cadherin (green) in section of P21 proximal small intestinal. Note that GORASP2 staining is no longer visible upon TAM injection at P1. **B:** Visualization of GORASP2 (red) and E-cadherin (green) in small intestine budding organoids (confocal section). Note that GORASP2 staining is no longer visible upon 4 OT incubation for 7 days. **C:** Western blot showing that GORASP2 is no longer expressed in organoids treated with 4 OT as in B. E-cadherin is the loading control.

Taken together, we conclude that TAM/4 OT treatment of cDKO mice/cDKO organoids, results in the generation of a double KO GORASP1 and 2(DKO mice/DKO organoids).

### No visible loss of Golgi stacking in DKO tissue

Small intestine (cDKO) and double knockout (DKO) were processed for conventional electron microscopy for an extensive comparative analysis of the organization of the Golgi. At P10(*Supp fig. S2A, B*; Figure 3C), P21(Figure 3A), P42(*Supp fig. S2A, B*; Figure 3C), the Golgi stacks in the DKO small intestines were prominently visible at each time point. The number of cisternae per stack was largely unchanged, averaging around 3 cisternae per stack (mean ±SD: P10 cDKO: 3,24±0,59/DKO: 2,71±0,61; P42 cDKO: 3,09±0,52/DKO 3,15±0,46).

**Figure 3:**
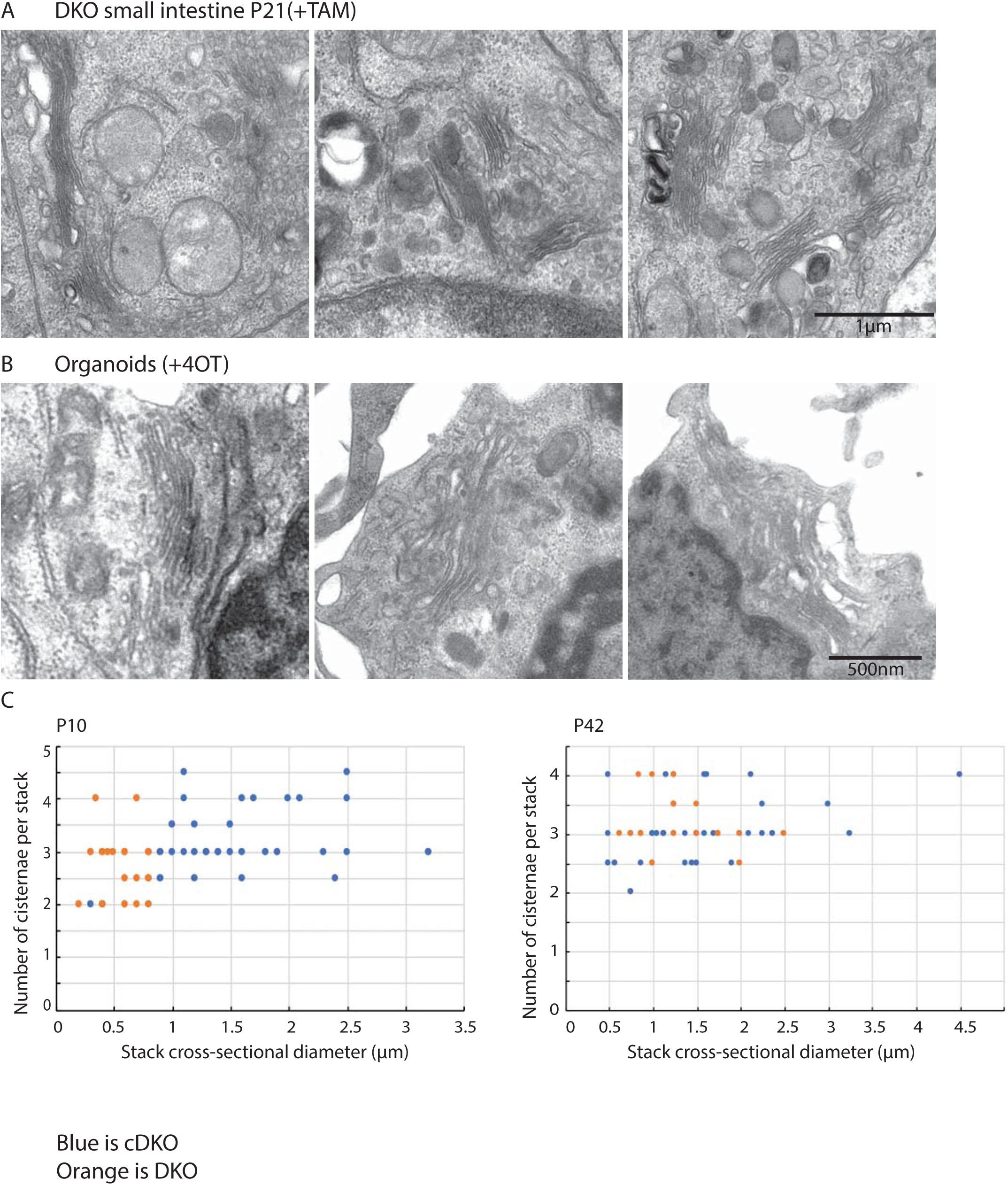
Double KO of GORASP 1 and 2 does not lead to Golgi unstacking. **A:** Electron micrograph of Golgi stacks in P21 small intestine from mouse treated with TAM at P1. Note that the Golgi are still stacked. **B:** Electron micrograph of Golgi stacks in small intestine organoids treated with 4 OT for 7 days. Note that the Golgi are still stacked. **C:** Quantification of the stack morphometry plotting the stack cross-sectional diameter versus the number of cisternae per stack in the small intestine of mice (±TAM at P1) at P10, P42. Blue is the cDKO and orange the DKO.Note that the cross-sectional diameter is reduced upon TAM/4 OT treatment but that the number of cisternae per stack is largely unchanged.

However, we noticed a quantitative change in the dimensions of the stacked cisternae. The cross-sectional diameters were shorter by a range of an eighth to two thirds (mean diameter μm ±SD: P10DKO: 0,58±0,27; P42 DKO 1,32±0,50) compared to the cDKO (mean diameter μm ±SD: P10cDKO: 1,52±0,63; P42 cDKO: 1,50±0,80). Of note, cells present in the stroma also exhibited Golgi stacks, in both the cDKO and DKO mice. Overall, these results show that GORASP1 and 2 are not required to stack the core of the Golgi cisternae in mouse small intestine, but they show clear functions on peripheral regions of the cisternae, which manifest as shorter than their control counterpart.

The rims of Golgi cisternae are highly fenestrated and It has been reported that conditions that exaggerate COPI vesicle formation, such as treatment with GTPJS or incubation of isolated Golgi membranes with mitotic cytosol, convert cisternal rims into small vesicles without severely affecting the core that remain stacked (Warren, 1985). This suggests that the rims of the cisternae are functionally different from the cisternae core. This also suggests the possibility that the core is stacked by a mechanism different from that linking the rims in a Golgi stack. GORASPs have the property of linking surfaces and bind tethers, golgins and SNAREs, components needed for COPI vesicle fusion (Ahat, Li, & Wang, 2019; Wei & Seemann, 2017). Perhaps, this linking controls vesicle budding/fusion at the rims. Unlinking of cisternae laterally by GORASP removal would expose more cisternal membrane to COPI vesicle budding, more than enough to compensate for the diminished fusion since the needed components are no longer presented properly (by GORASPs). This would create a new steady-state with more local COPI vesicles driving increased flux. The increased local vesicles would "contain" the rim membrane and explain the decreased cisternal diameter. In other words, GORASPs control the linking of cisternal rims laterally into a Golgi ribbon whereas another hitherto uncharacterized set of components additionally cement the core of Golgi cisternae.

### GM130 does not compensate for the loss of GORASPs

The cis Golgi associated, peripheral protein GM130 is diffusely dispersed around the Golgi in the double GORASP CRISPR KO HeLa cells (see, Figure 5A in (Bekier et al., 2017)). This could potentially compromise the Golgi stacking caused by the GORASP double KO. In other words, the stacking we observe the GORASP1 and GORASP2 DKO small intestine and organoids could be due to the presence of GM130, or even its upregulation to compensate for the loss of GORASPs. Therefore, we stained the same tissue sections as above for GM130, and found that while GM130 was abundant in the cDKO material, it was completely absent in the DKO tissue (Figure 4A). This strongly indicates that GM130 is degraded in GORASP1 and 2 lacking cells. Interestingly, it has been shown earlier that depletion of GORASPs by CRISPR resulted in reduction of GM130 by 75% (Bekier et al., 2017). The severity of the effect on GM130 levels might depend on the levels of GORASPs and merits further investigation.

**Figure 4:**
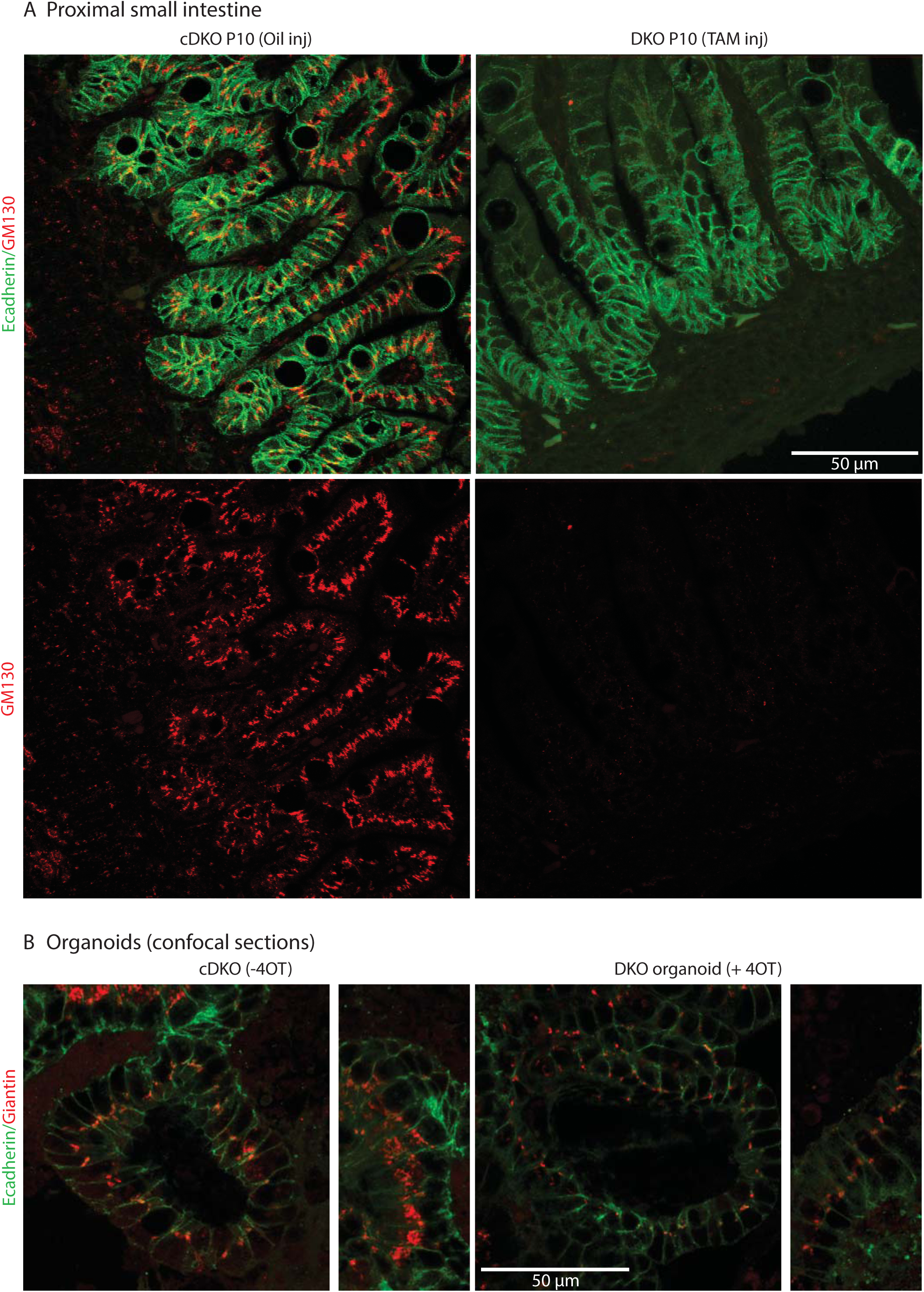
GM130 is degraded in GORASPs knockout cells and does not affect cisternal stacking. **A:** Visualization of GM130(red) and E-cadherin (green) in section of P10 proximal small intestine in mice injected with either oil or TAM at P1. Note that GM130 staining is no longer visible upon TAM injection at P1. **B:** Visualization of Giantin (red) and E-cadherin (green) in budding organoids treated or not with 4 OT (confocal section). Note that the Giantin staining is still prominent in every cell after 4 OT although the intensity of staining is decreased, possibly reflecting the Golgi ribbon unlinking and dispersion of the smaller stacks in the cell cytoplasm.

## Conclusions

Our data provide a unifying framework for GORASP function. We suggest that a Golgi cisternae is composed of at least two discrete functional domains: a core that can stack in a GORASP-independent manner, and rims that are scaffolded and linked laterally by GORASPs (Ladinsky, Mastronarde, McIntosh, Howell, & Staehelin, 1999; Rabouille & Linstedt, 2016). Sub-cisternal localization of Golgi components indicate specificity in location of Golgi components associated with the core or the periphery of a cisternae (Kweon et al., 2004; Tie, Ludwig, Sandin, & Lu, 2018). GORASP proteins are present at higher levels at the rim of a cisternae than the core (Tie et al., 2018), which fits well with the function of GORASPs at the rims of the Golgi cisternae. Depletion of GORASPs alters the physical and functional properties of the rims and promotes hyper-vesiculation, making for a smaller Golgi apparatus. Interestingly, this increased vesiculation leaves the core of the Golgi largely intact, while compounds like Ilimaquinone, completely vesiculate Golgi stacks (Takizawa et al., 1993). These data indicate that there are different mechanisms to control Golgi cisternae-associated adhesion and vesicle formation at the core vs at the rims. GORASPs affect one, while ilimaquinone acts by inactivating both mechanisms of cisternal adhesions.

Genetic data show that deletion of the single GORASP gene in *Dictyostelium discoideum* and yeast, inhibits starvation-specific secretion of signal sequence lacking ACBA/Acb1(Duran et al., 2008; Kinseth et al., 2007; Manjithaya, Anjard, Loomis, & Subramani, 2010). In Drosophila, depletion of the single GRASP ortholog prevents a Golgi-bypassed delivery of the transmembrane protein Integrin, to the plasma membrane (Schotman, Karhinen, & Rabouille, 2008). Clearly, genetics tells us that GORASPs have a role in unconventional secretion. This conclusion is now supported by a large number of reports showing the involvement of GORASPs in other systems including mammalian cells (Feinstein & Linstedt, 2008; Noh et al., 2018; van Ziel, Largo-Barrientos, Wolzak, Verhage, & Scheper, 2019; Wu, Li, & Zhao, 2020). Although transport by COPII and COPI is not required for unconventional secretion, the essential role of GORASPs in this process suggests a link to the Golgi membranes. One possibility is that the rims of Golgi cisternae are a necessary source of membranes or a surface for proteins required for unconventional secretion. These two roles in rim stacking and unconventional secretion are not mutually exclusive, but likely represent the use of GORASPs to stabilise and control membrane flux and vesicle biogenesis at fenestrated membranes, as are seen at cisternal rims of the Golgi apparatus or CUPS (compartment for unconventional secretion). These could be COPI vesicles during conventional secretion and COPI-independent vesicles to transport specific components for unconventional protein secretion to CUPS. The latter would be functionally analogous to the export of Atg9 by small vesicles from late Golgi to create an autophagosome.

## Materials and Methods

### Mouse genetics

The generation and characterization of GORASP1 knock-in (KI, that act as a KO) is described in (Veenendaal et al., 2014). Briefly, GORASP2 KI/KO mice were generated at McLaughlin Research Institute (Great Falls, Montana, US) using the EUCOMM/KOMP knockout first conditional-ready targeted ES cell resource (http://www.mousephenotype.org/about-ikmc/eucomm-proram/eucomm-targeting-strategies; targeted trap tm1 a allele). In short, Gorasp2 tm1 a(EUCOMM)Hmgu ES cells were injected into C57 BL/6 J-Tyr<c-2J> blastocysts to generate chimeras that were bred to C57 BL/6 J-Tyr<c-2J> mice. Offspring were subsequently brother-sister mated for several generations.

We first attempted to generate a double KI by crossing the mice to homozygosity, but we were unable to recover pups of the right genotype, suggesting that the loss of both GORASPs is critical for embryonic development. We therefore resorted to use the GORASP2 cFlox allele of the GORASP2 KI mouse: GORASP2 cFlox mouse. Conditional knock-out (cFlox) mice were generated by breeding the tm1 a allele carrying mice with Flp-recombinase expressing mice, removing the LacZ and neomycin cassettes and resulting in mice expressing endogenous GORASP2, with exon 5 being flanked by loxPsites (tm1 c allele) (Figure 1B). Null alleles were generated by breeding cKO mice with mice expressing Cre-recombinase under the control of Zp3 promoter, eliminating exon 5 in the female germline (tm1 d allele).

Primers 5’-ccgatcctttcctcagccgtcagc-3’, 5’-cctcaaggccctaccttagc-3’ and 5’- tgaactgatggcgagctcagacc-3’ were used to genotype the offspring using genomic DNA extracted from ear punches (Mouse Direct PCR kit, Biotools).

We then crossed the GORASP2 cFlox mouse and the GORASP1 KI mouse to a Rosa Cre-ERT2 mouse, and through back crossing, generated a GORASP1 KI HOM; GORASP2 cFlox HOM; Rosa Cre ERT2 HOM mouse that we called the “cDKO” line. Upon injection of 20 μl of 5 mg/ml of Tamoxifen (TAM, Sigma, T5648) 1 day after birth (Postnatal day 1, P1), GORASP2 exon 5 is excised (GORASP2 FLOX, Figure 1C, D) and the full-length protein is no longer expressed (Result section). Mice sacrificed at P42 were re-injected at P21 and P39 with 170 μl of 5 mg/ml of Tamoxifen. Controls were performed by injected oil to a parallel litter of the same genotype. Note that exon 6 to 10 are still present in the genomic locus. This yields a GORASP2 mRNA that misses exon 5 and a complete loss of GORASP2 protein (Immunofluorescence and Western blot) *(Supp fig. S1A-C).*

### cDKO Organoids

We generated small intestine organoids using the published procedure (Beumer et al., 2018) using a 6-weeks female GORASP1 KI HOM; GORASP2 cFlox HOM; Rosa Cre ERT2 HOM mouse. We used Cultrex RGF BME, type II (3533-005-02) instead of Matrigel. To deplete GORASP2 gene, 100 nM 4-hydroxy Tamoxifen (4 OT, Sigma, H7904) for 7 days was added on budding large organoids.

### Monitoring exon 5 excision

For DNA extraction, 20 mg of proximal gut tissue (flushed, washed and snap frozen) was used as described in DNeasy Blood & Tissue Kits (Qiagen, 69504) to recover 30 ng/μl DNA.

RNA was extracted from 650-700 organoids harvested with Corning Cell Recovery Solution (Corning, 354253) removing BME gel. After washing with PBS, the RNA was extracted as recommended by the RNeasy Mini Kit (Qiagen, 74104). This yielded about 20 ng/μl of RNA. Of all samples, a total of 75 ng cDNA in 20μl was synthesized by GoScript Reverse Transcriptase (Promega, A5000).

### Primers for PCR shown in Figure 1 and Supp fig. S1

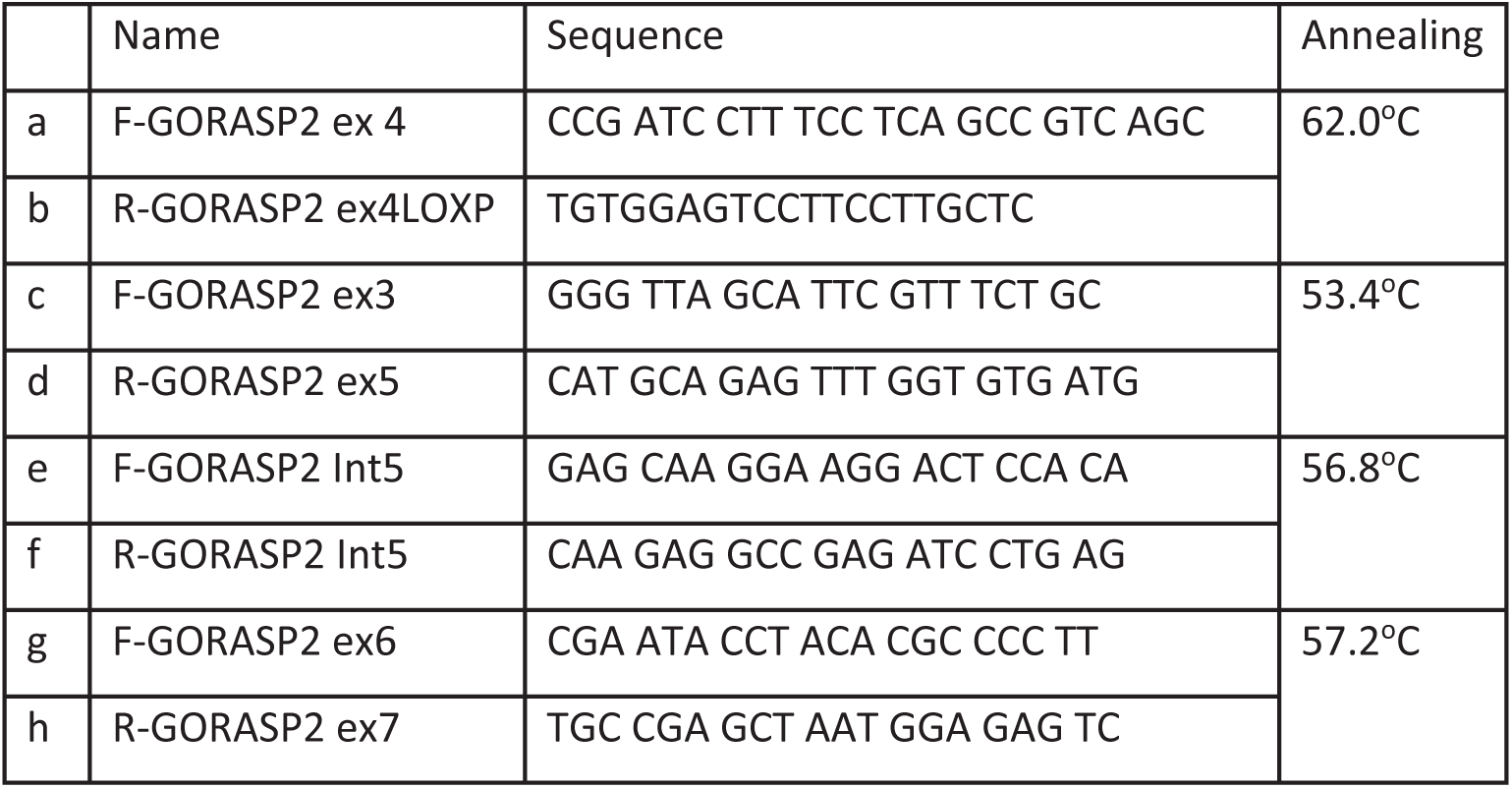
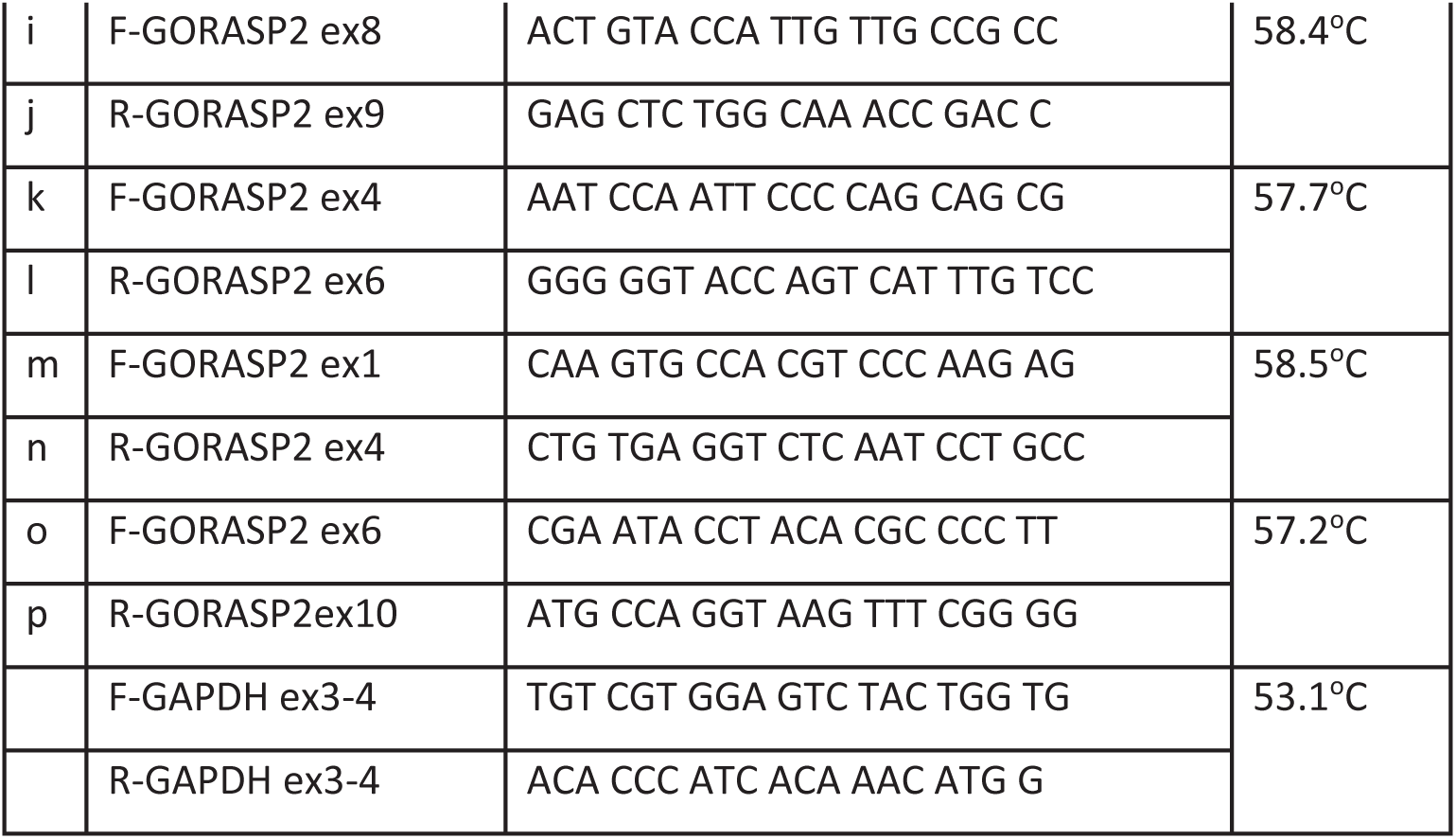

### Western Blot

1500 organoids treated or not with 100 nM 4 OT for 7 days were harvested using 1 ml Corning Cell Recovery Solution (Corning, 354253) per ∼ 60 μl of BME gel. After centrifugation, they were lysed in 100μl RIPA buffer (50 mM Tris-HCl at pH 7.5, 150 mM NaCl, 1% NP-40, 0.5% Na-deoxycholate, 0.1% SDS, 5 mM EDTA, 50 mM NaF, 1 mM PMSF supplemented with Roche 1X Halt Protease inhibitor cocktail), incubated for 1 hour at 4 °C, and sonicated using a Bioruptor PLUS (Diagenode) for 30 cycles of 30 seconds spaced by 30 second of cooling down. Total protein concentration in lysates was determined using Pierce BCA Protein Assay Kit (Thermo Scientific, 23227). The protein concentration in the lysates was 1.74μg/μl. Proteins were fractionated in 10% polyacrylamide gel and blotted on PVDF membrane. Milk was used as a blocking agent.

Membrane was incubated with a Rabbit anti GORASP2(Rich, against aa 232–454, gift from M. Bekier, (Xiang & Wang, 2010) at dilution 1/1000), and a Mouse monoclonal Antibody to E-cadherin (BD Biosciences 610182 at 1/5000).

The visualization was performed after incubation with secondary antibodies coupled to HRP (GE Healthcare, NA931 and NA934) and Clarity Western ECL Substrate (BioRad) using ImageQuant LAS4000 ECL (GE Healthcare).

### Immunofluorescence microscopy

The small intestine of oil and TAM injected cDKO mice were dissected at P10, P21 and P42, flushed, fixed in formalin (Merck) overnight. It was then processed and embedded in paraffin.

Sections were cut on a microtome, rehydrated, boiled in citrate, blocked with 1% BSA and stained with primary antibodies followed by fluorescent secondary antibodies as described (Beumer et al., 2018). Slides were observed under a SPE Leica microscope.

Budding organoids treated with and without 100 nM 4 OT for 7 days and were fixed directly in the gel leading them to stick tightly to the glass coverslip that is set underneath the gel drop. Coverslips were simply removed and processed for IF after permeabilization using 0.5% triton X-100 for 30 min.

We used a Rabbit anti GORASP2 antibody (Rich, against aa 232–454) from M. Bekier, (Xiang & Wang, 2010) at dilution 1/200); a rabbit anti GM130 antibody (MLO8, gift from M. Lowe at dilution 1/100); a mouse anti E-cadherin antibody (BD Biosciences 610182 at 1/1000). Primary antibodies were coupled to donkey anti Rabbit Alexa 568(Invitrogen, A10042) or goat anti mouse Alexa 488(Invitrogen, A-11001).

### Electron microscopy

A small part of the intestine that is close to the part processed for immunofluorescence was fixed in Karnovsky (2% Glutaraldehyde, 2% paraformaldehyde, 0.25 mM CaCl2, 0.5 mM MgCl2, in phosphate buffer (0.1M at pH7.4)) overnight at room temperature followed by 1% PFA at 4 oC. After processing, they were embedded in epon resin. Ultrathin plastic sections were cut and examined under a Jeol microscope (Slot & Geuze, 2007). Images of crypts and villi were acquired and stitched together to allow a view of the small intestine section both at low and a high magnification.

### Quantification

The cross-sectional diameter of each stack that is visible in the intestinal section was measured using a ruler on high magnification pictures (20,000 x). Using the same pictures, the number of cisternae per stack was counted and plotted again the cross-sectional diameter.

Each recognizable Golgi area is accounted for, whether a stack is present or not. 0 in the graph means that a Golgi area was identified but that no stacks were present.

---

**Fig. S1:**
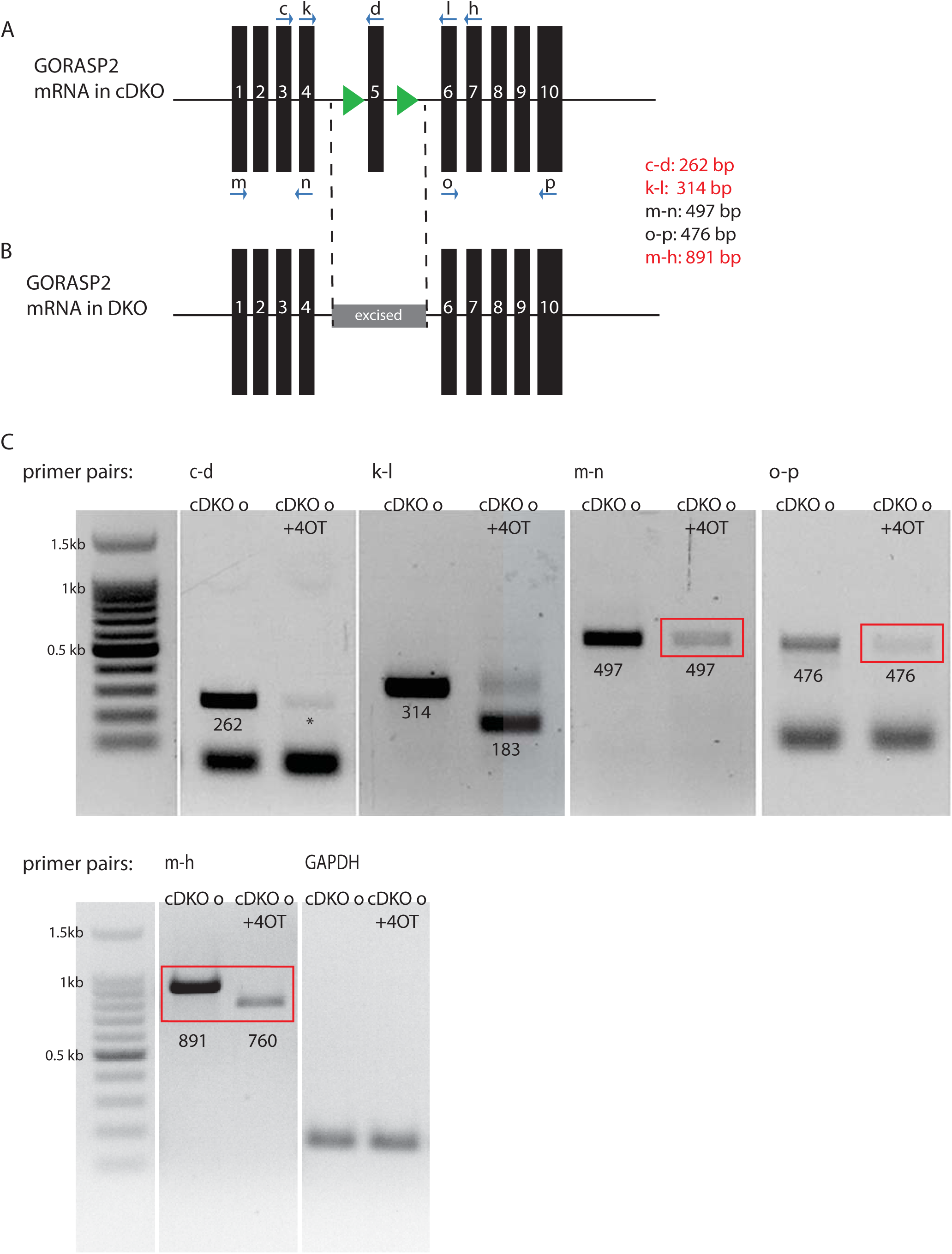
GORASP2 mRNA in cDKO and DKO organoids. **A:** Schematic of the GORASP2 cFlox mRNA. The exon 5 is now flanked by two LOXP sites. **B:** Schematic of the GORASP2 FLOX allele upon 4 OT treatment. The exon 5 is excised. **C:** A series of PCR using different combination of primers showing that exon 5 is excised upon 4 OT inhibition (c-d) and that the sequence between exon 4 and 6 is shorter (from 314 bp to 183 bp, k-l), that exon 1-4 is produced at a lower amount than control (red box, m-n) and that exon 6 to 10 is also expressed at a lower amount than control (red box, o-p). Exon 1-7 produced both a lower amount and a shorter sequence compared to the control.

**Fig. S2:**
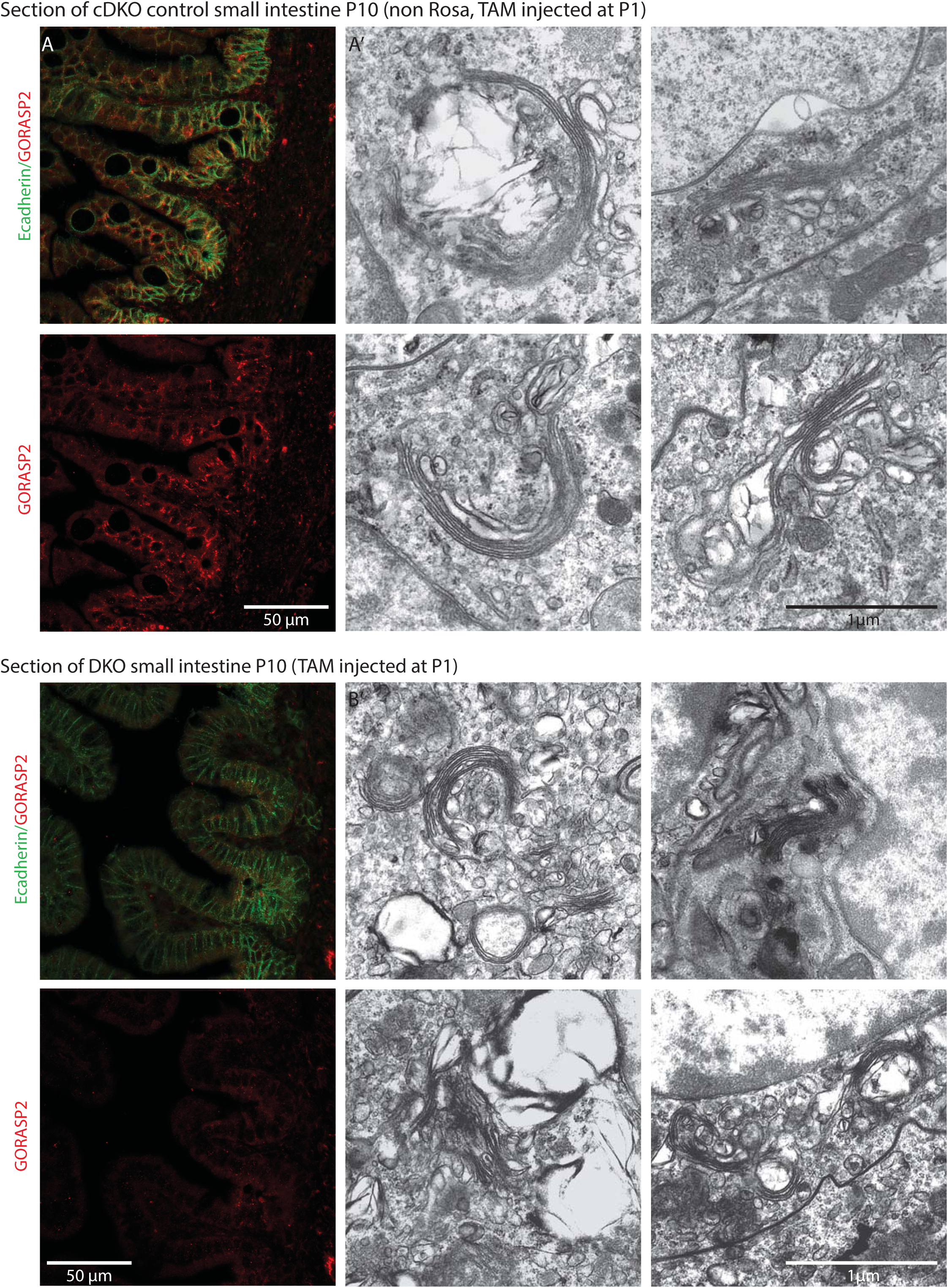
Double KO of GORASP 1 and 2 does not lead to cisternal unstacking in P10 mouse small intestine. **A, A’:** Visualization of GORASP2(red) and E-cadherin (green) in section of P10 proximal small intestine of control mice injected with oil (A), and corresponding Golgi stack profiles (A’). Note that the stacks are large and contain 2-4 cisternae. **B, B’:** Visualization of GORASP2(red) and E-cadherin (green) in section of P10 proximal small intestine of control mice injected with TAM at P1(B), and corresponding Golgi stack profiles (B’). Note that GORASP2 staining is no longer visible upon TAM injection, yet the Golgi stacks are clearly visible. They are shorter but contain the same number of cisternae. Quantification is shown in Figure 3C.

**Fig. S3:**
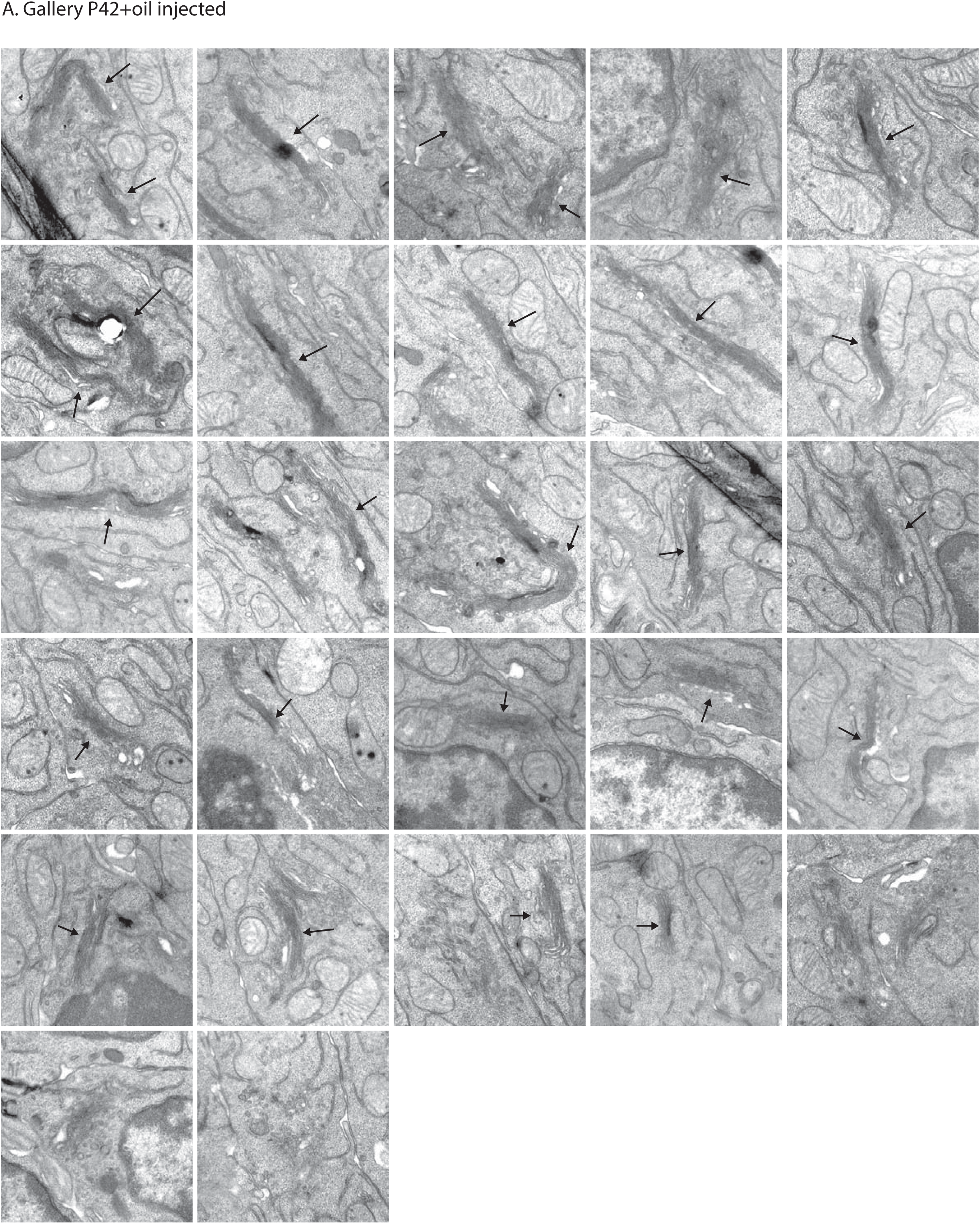
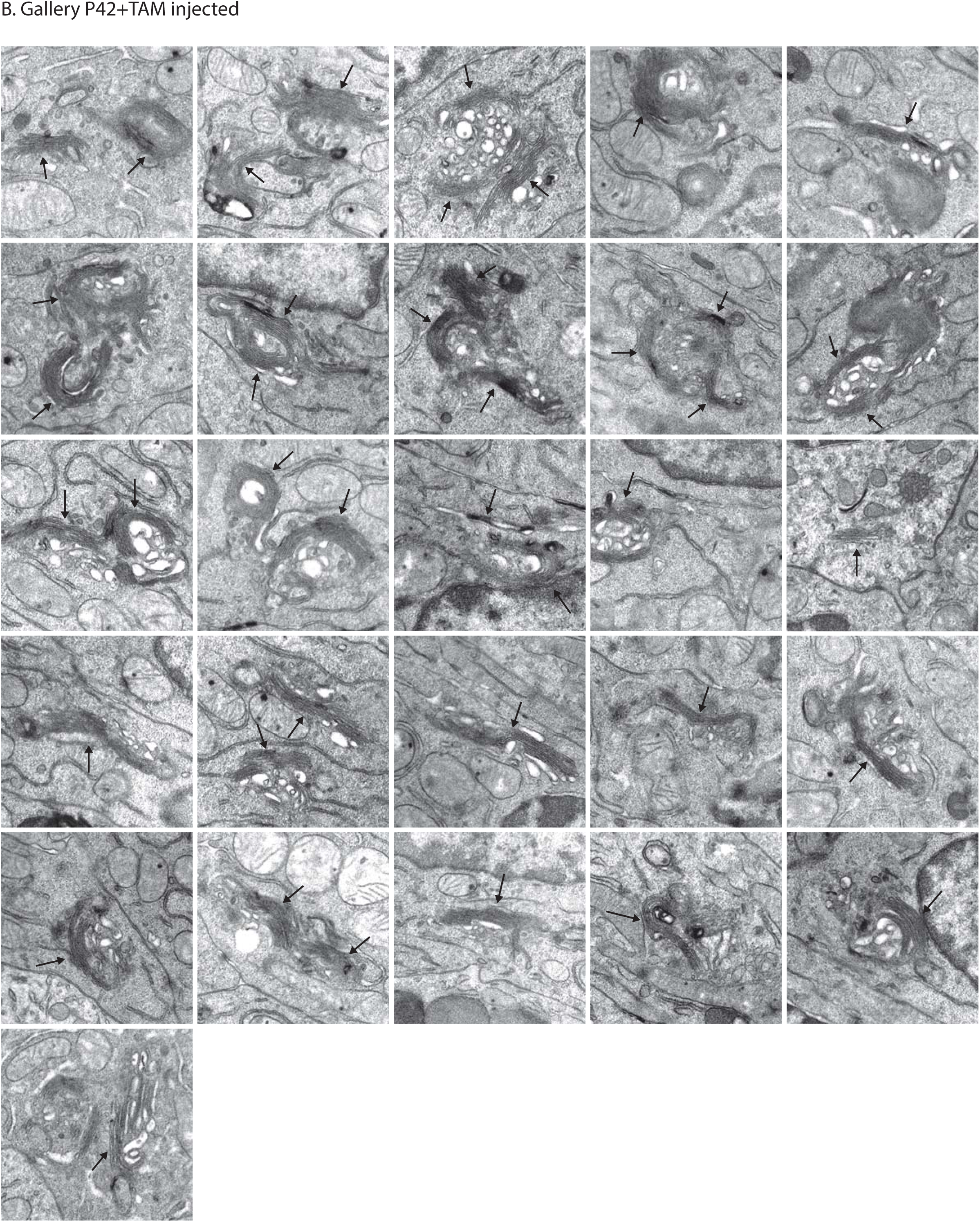
Double KO of GORASP 1 and 2 does not lead to Golgi unstacking in P42 mouse small intestine. **A:** A gallery of Golgi stack profile in sections of small intestine control mouse at P42 injected with oil at P1. **B:** A gallery of Golgi stack profile in sections of small intestine mouse at P42 injected with TAM at P1. Note that as in P10, the stacks are shorter in cross-sectional diameter but that the number of cisternae per stack is unchanged upon TAM injection (Quantification is shown in Figure 3C).

## Acknowledgements

We thank Graham Warren for helping us write a balanced discussion. VM is an Institució Catalana de Recerca i Estudis Avançats professor at the Centre for Genomic Regulation, and the work in his laboratory is funded by grants from the Ministerio de Economía y Competitividad (MINECO) (SEV-2012-0208, BFU2013-44188-P, CSD2009-00016)

## Notes

### Competing Interest Statement

The authors have declared no competing interest.

